# Alfaxalone activates Human Pregnane-X Receptors with greater efficacy than Allopregnanolone: an in-vitro study with implications for neuroprotection during anesthesia

**DOI:** 10.1101/2020.09.05.284075

**Authors:** Juliet.M. Serrao, Colin.S. Goodchild

**Affiliations:** Drawbridge Pharmaceuticals Pty Ltd, Malvern, Victoria, Australia

## Abstract

**Background:** Alfaxalone is a fast acting intravenous anesthetic with high therapeutic index. It is an analogue of the naturally-occurring neurosteroid, allopregnanolone which has been implicated in causing neuroprotection, neurogenesis and preservation of cognition, through activation of pregnane X receptors in the central nervous system. This study investigated whether alfaxalone can activate human pregnane X receptors (h-PXR) as effectively as allopregnanolone.

**Methods:** Allopregnanolone and alfaxalone were dissolved in dimethyl sulfoxide to make allopregnanolone and alfaxalone treatment solutions (serial 3-fold dilution concentration range, 50,000 – 206 nM). Activation of h-PXR by these ligand solutions compared with vehicle control was measured by an in-vitro method using human embryonic kidney cells (HEK293) expressing h-PXR hybridised and linked to the firefly luciferase gene. Ligand binding with and activation of h-PXR in those cells caused downstream changes in luciferase activity and light emission. That activity was measured as relative light units using a plate-reading luminometer, thus quantifying the changes in h-PXR activity caused by the ligand applied to the HEK293 cells. Ligand log concentration response curves were constructed to compare efficacy and potency of allopregnanolone and alfaxalone.

**Results:** Allopregnanolone and alfaxalone both activated the h-PXR to cause dose-related light emission by the linked firefly luciferase. Control solutions (0.1% dimethyl sulfoxide) produced low level light emissions. Equimolar concentrations of alfaxalone were more efficacious in activation of h-PXR: 50,000 nM, p = 0.0019; 16,700 nM, p = 0.0472; 5,600 nM, p = 0.0031 [Brown-Forsythe and Welch ANOVA].

**Conclusions:** Alfaxalone activates human-pregnane X receptors with greater efficacy compared with the endogenous ligand allopregnanolone. These results suggest that alfaxalone sedation and anesthesia may be accompanied by beneficial effects normally caused by the physiological effects of allopregnanolone, namely neuroprotection, neurogenesis, and preservation of cognition.

## Introduction

Pregnane X Receptor (PXR; NR112) is a nuclear receptor that binds with, and is activated by, a variety of xenobiotic and endogenous compounds, including naturally occurring steroids, pregnenolone, progesterone and allopregnanolone ^1;2^. Allopregnanolone is a metabolite of progesterone synthesized in the central nervous system where it promotes neurogenesis and neuroplasticity ^3;4^. PXR activation by allopregnanolone has also been linked to neuroprotection ^5^. Further, these properties are due to PXR activation by allopregnanolone stimulating the production of brain-derived neurotrophic factor (BDNF) ^5–8^. The most important functions of BDNF include: regulation of neuro-, glio-, and synapto-genesis; neuroprotection; and control of short- and long-lasting synaptic interactions responsible for memory and cognition ^9^.

Alfaxalone, (3α-hydroxy-5α-pregnane-11,20-dione) is a pregnane steroid which has potent anesthetic and sedative properties by actions at gamma aminobutyric acid type A (GABAA) receptors ^10^. It is an allopregnanolone and progesterone analogue (Figure 1), but it is devoid of conventional progestogen or endocrine hormonal activity ^11^. An aqueous formulation of alfaxalone (Phaxan™) has been developed for use as an intravenous sedative and anesthetic ^12;13^. This development occurs at a time when most commonly-used anesthetics are under investigation for neurotoxic effects ^14–16^. Alfaxalone differs from those commonly used anesthetics in that it is a structural analogue of a naturally occurring neuroprotective hormone, allopregnanolone. It is unknown whether alfaxalone can activate human pregnane X receptors (h-PXR) although its structural similarity to allopregnanolone suggests that it may do so. If it is found that alfaxalone can activate mechanisms utilized by allopregnanolone to cause neuroprotection, e.g., PXR, there would be implications for the use of alfaxalone in clinical anesthetic practice. This study set out to compare the activation of h-PXR in-vitro by equimolar concentrations of allopregnanolone and alfaxalone.

**Figure 1:**
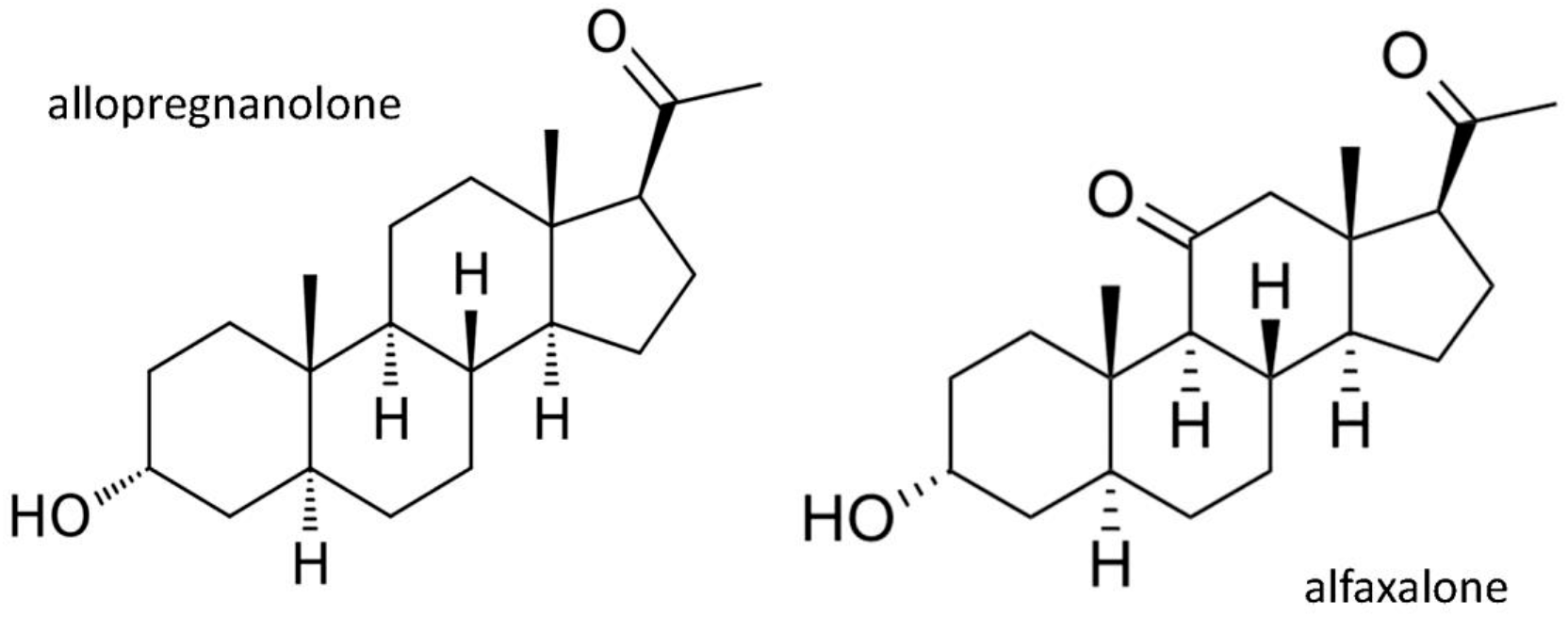
Allopregnanolone and alfaxalone: molecular structures compared

## Materials and methods

### Assay Platform

The activation of h-PXR by allopregnanolone and alfaxalone was measured using an in-vitro preparation by INDIGO Biosciences, Inc., 1981 Pine Hall Road, State College, PA, USA. This consisted of suspensions of “reporter cells” in multi well plates; human cells (HEK293) expressing the nuclear receptor, h-PXR (NR1I2). These cells were engineered to express a hybrid h-PXR, in which the N-terminal sequence encoding the binding domain was substituted with yeast GAL4-binding domain. The rest of the native h-PXR, i.e., the ligand binding and C-terminal domains, remained intact and functional. Ligand (e.g., allopregnanolone) interaction with the receptor, caused it to bind to the GAL4 DNA binding sequence. The latter was functionally linked to firefly luciferase reporter gene. Binding with, and activation of the h-PXR, caused downstream changes in luciferase activity and light emission from the treated reporter cells. That luminescence was measured as relative light units (RLU) using a plate-reading luminometer, thus measuring the changes in h-PXR activity caused by the ligand applied to the HEK293 cells in each well plate.

### Test Compounds

The test compounds alfaxalone and allopregnanolone were obtained from Sigma-Aldrich (400 Summit Drive, Burlington, MA 0180).

### Assay Methods

A suspension of reporter cells was prepared in cell recovery medium. Stock solutions of allopregnanolone and alfaxalone were diluted in dimethyl sulfoxide (DMSO) to generate concentrated stock solutions (1000 x the highest test concentration). These intermediate stocks were diluted directly into compound screening medium containing 10% charcoal-stripped foetal bovine serum to generate allopregnanolone and alfaxalone treatment solutions with concentrations at double the final planned testing levels: 100,000; 33,333; 11,111; 3,704, 1,235; and 412 nM respectively. 100 μL of each prepared treatment solution was dispensed into quadruplicate assay wells pre-filled with a 100 μL suspension of reporter cells, thereby achieving the desired final treatment concentrations as shown in Table 1. The concentration of residual DMSO in all assay wells was 0.1%.

**Table 1.**
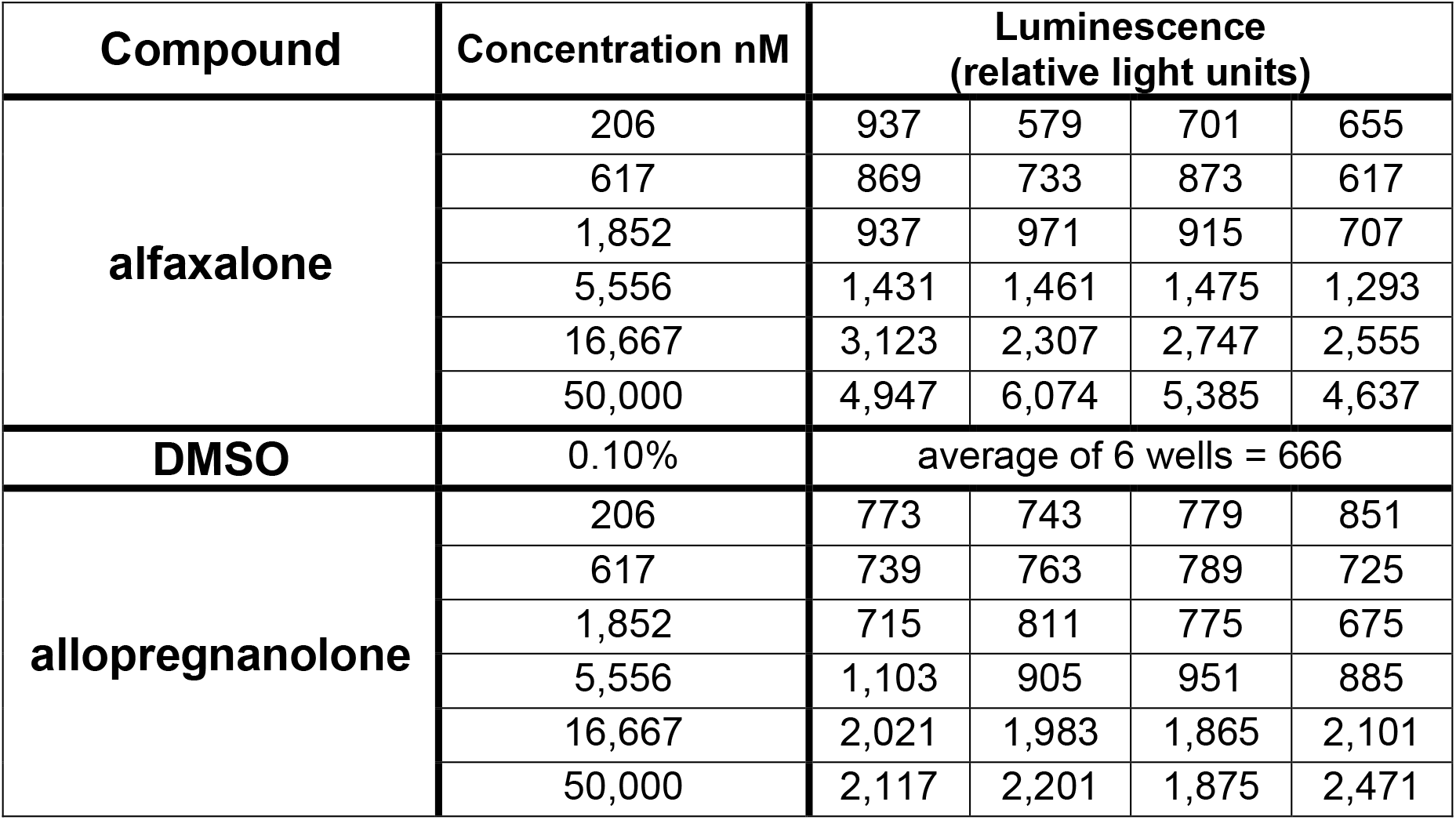
hPXR Assay Results: The luminescence (expressed as relative light units, RLU) caused by a range of concentrations of allopregnanolone and alfaxalone; four wells at each concentration. Background luminescence of 0.1% solution of dimethyl sulfoxide (DMSO), the solution used to dissolve the neurosteroids, is also shown.

Assay plates were incubated for 22-24 h in a cell culture incubator (37°C / 5% CO2 / 85% humidity). Following the incubation period, the treatment media were discarded and 100 μL of luciferase detection reagent (Indigo Biosciences) was added to each well. The resulting luminescence (relative light units; RLUs) was measured in each well using a luminometer. The recording from each well was entered into an Excel spreadsheet. Agonist concentration/activity response curves were plotted with non-linear curve fitting, using GraphPad Prism software (version 8.4.1; GraphPad Software, 2365 Northside Dr. Suite 560, San Diego, CA 92108).

## Results

Allopregnanolone and alfaxalone both produced aqueous solutions up to maximum concentration of 50,000 nM in 0.1% DMSO. Higher concentrations of drug were not possible because the higher concentrations of DMSO necessary to achieve drug dissolution disrupt normal functions of the HEK reporter cells. Both pregnane steroids activated the hybrid h-PXR to cause dose-related light production by the linked firefly luciferase compared with control (DMSO) solutions (table 1 and figure 2).

**Figure 2.**
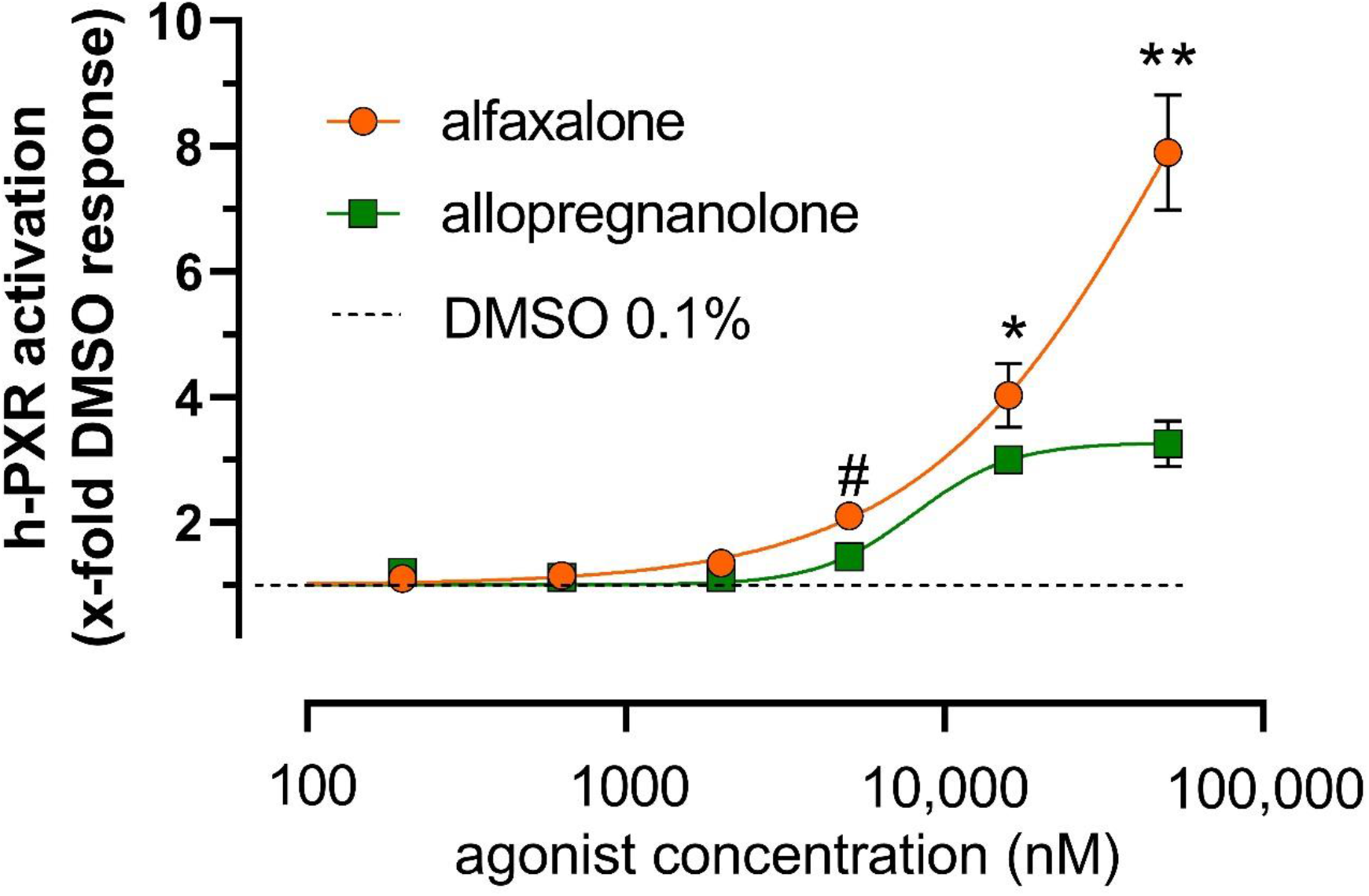
Ligand log concentration dose response curves: hPXR activation. Luminescence caused by the ligand interaction with h-PXR in each well was expressed as a multiple of the average luminescence caused by 0.1% DMSO vehicle controls (n=6). Points shown are means (n=4) and bars ± 95% confidence intervals (not shown if smaller than the symbol size). Statistical comparison using Brown-Forsythe and Welch ANOVA (GraphPad Prism version 8.4.1) confirmed alfaxalone is more efficacious than allopregnanolone in activating h-PXR at: 50,000 nM [**, p=0.0019]; 16,700 nM [*, p = 0.0472]; and 5,600 nM [#, p = 0.0031].

Demonstration of the maximum dose response for alfaxalone was not possible because of the inability to prepare more concentrated solutions with DMSO. Analysis of the log10 ligand concentration response relationships (Figure 2; GraphPad Prism 8 software) revealed that the allopregnanolone concentration that caused 50% of maximum effect (EC50) was 9192, 6716 – 12581 nM (mean, 95% CI); a comparable figure for alfaxalone could not be calculated because the maximal response was not achieved.

However, the maximum values at the top of the log concentration response curves, expressed as x-fold activation of h-PXR (control DMSO = 1) by 50,000 nM solutions were very different for the two ligands: 3.25, 2.66 – 3.83 (allopregnanolone mean, 95% CI); 7.90, 6.44 – 9.36 (alfaxalone mean, 95% CI) [p=0.0019; Brown-Forsythe and Welch ANOVA – figure 2]. Furthermore, h-PXR activation by lower intermediate concentrations of the two agonists were also significantly different: 16,700 nM, p = 0.0472; 5,600 nM, p = 0.0031 [Figure 2; Brown-Forsythe and Welch ANOVA], alfaxalone being the more efficacious compound in activating h-PXR.

## Discussion

The results of the study reported here show that alfaxalone does bind with and activates h-PXR and further that it is a more efficacious ligand than allopregnanolone in activating h-PXR in this model. Conclusions about the relative potency of the two pregnane ligands were not possible because the plateau portion of the alfaxalone log10 concentration response curve was not demonstrated. This was due to the inability to dissolve alfaxalone in water using 0.1% DMSO at concentrations higher than 50,000 nM. This result suggests that sedation and anesthesia achieved using alfaxalone may be accompanied by the effects caused by interaction of allopregnanolone with brain PXR such as stimulating the production of BDNF ^5–8^, which has been shown to have important roles in regulation of neurogenesis, gliogenesis, and synaptogenesis, as well as in neuroprotection, and control of short- and long-lasting synaptic interactions that determine memory and cognition ^9;17^.

See Figure 3 for a schematic representation of the relationships between neurosteroids, PXR and BDNF and their effects on brain and neuronal function under conditions of anesthesia, stress, trauma, and inflammation. PXR activation by alfaxalone may also inhibit microglial hyperinflammatory responses in the central nervous system via neuroimmune regulatory proteins, such as CD55 ^18;19^. Glial-mediated inflammation in the CNS is caused by many factors such as trauma, hypoxia, stress, and also by β-amyloid in Alzheimer’s disease, all of which are common co-morbidities that complicate surgery and lead to poorer postoperative cognitive function ^19–21^. Glial-mediated inflammation is also a cause of deficits in cognition after bacterial and viral infections, such as COVID-19 ^22^. Alfaxalone sedation and anesthesia with simultaneous activation of PXR as described above clearly has the potential for improved CNS recovery.

**Figure 3.**
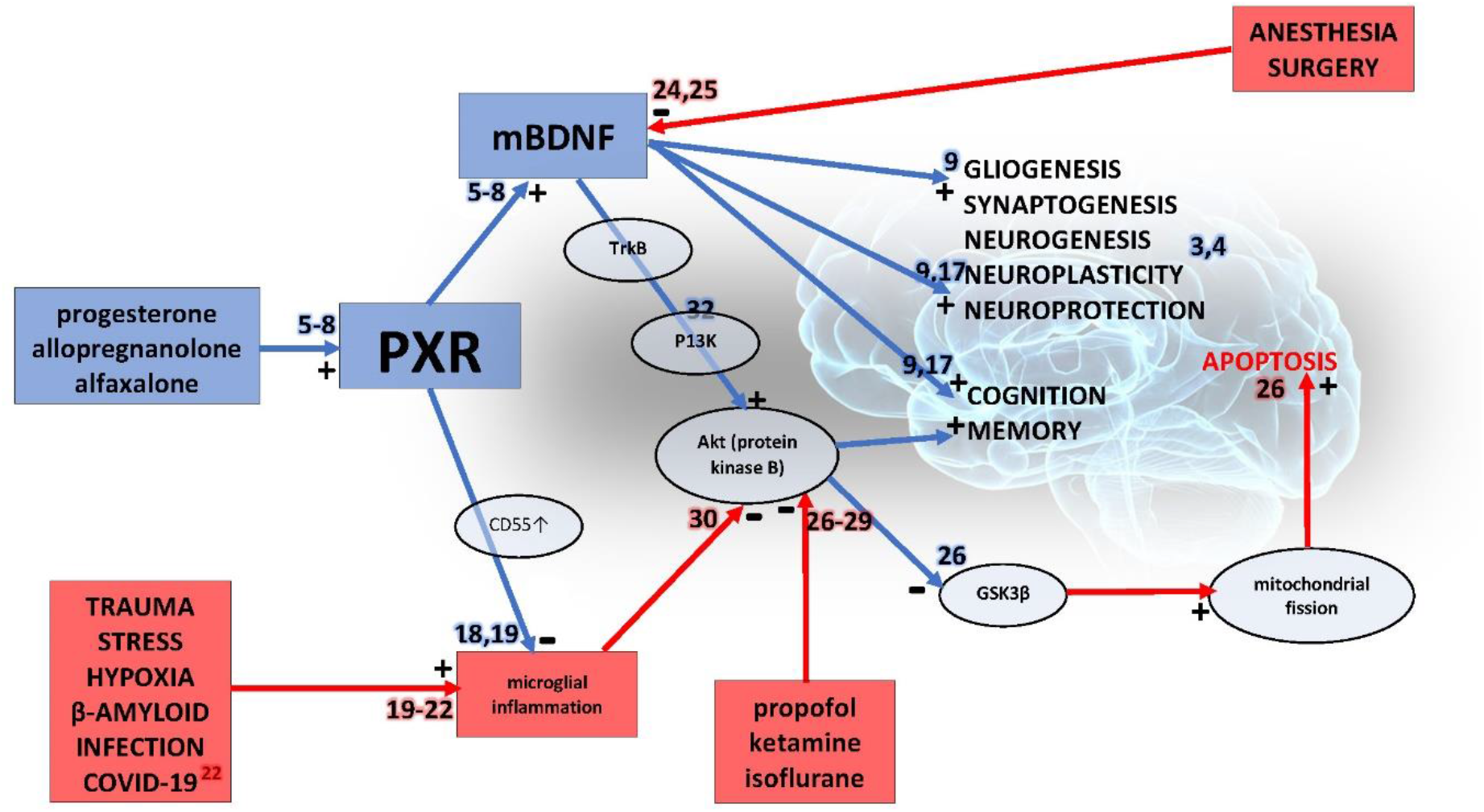
A schematic representation of the relationships between neurosteroids, PXR and BDNF and their effects on brain and neuronal function under conditions of anesthesia, stress, trauma, and inflammation. Neurosteroids progesterone, allopregnanolone and the neuroactive steroid alfaxalone bind with and activate pregnane X receptors (PXR) to cause increased production of mature brain derived neurotrophic factor (mBDNF) leading to neuroprotection and preservation of cognition in the face of trauma, stress, infection and the presence of anesthetics. Arrows denote a causal relationship, red for deleterious and blue for beneficial effects on brain function. A “+” or “-“ next to an arrow indicates a positive effect or negative effect on the target of the arrow. Small numbers adjacent to arrows refer to papers in the reference list, red background denoting deleterious effects and blue denoting beneficial effects

Alfaxalone is being developed for sedation and anesthesia in humans ^12;13^. This occurs at a time of controversy in anesthesia and critical care. Many preclinical studies published in the last two decades have reported neuronal damage and long-lasting cognitive impairment after administration of anesthetics, particularly to neonates and juveniles ^23^. It is debated whether these effects translate into human anesthesia practice and the root causes of the effects are controversial; neurotoxicity of the anesthetic drugs per se or neuronal damage and dysfunction caused by surgery, stress, infection, hypoxia, etc. as discussed above. However, it is true to say that all current commonly used anesthetic drugs have been reported to be neurotoxic in young animals. The United States Food and Drug Administration has issued an advisory warning concerning the possible dangers of administering those drugs in late-stage pregnancy and to babies and young children [https://www.fda.gov/drugs/drug-safety-and-availability/fda-drug-safety-communication-fda-review-results-new-warnings-about-using-general-anesthetics-and]. At the other end of the age spectrum, anesthetic drugs and surgery have been linked with acute delirium and long-term deficits in cognition in older persons ^24^. These have been linked to low levels of BDNF ^25^.

There are reports citing precise mechanisms for neurotoxicity of anesthetic drugs per se. Liu et al showed that propofol causes dose-dependent neuronal cell death at clinically relevant concentrations through a mitochondrial pathway; Akt (protein kinase B)/glycogen synthase kinase-3 (GSK3) ^26^. See Figure 3. This mechanism for neurotoxicity is also activated by ketamine and inhalational agents such as isoflurane ^27–29^. Further, the apoptosis caused by persistent neuroinflammation has been shown to involve the same Akt pathway as shown in Figure 3 ^30^. Sevoflurane anesthesia has been shown to induce an increase in the levels of pro-inflammatory cytokines in microglia, while decreasing activation of the phosphatidylinositol 3-kinase/protein kinase B (PI3K/Akt) pathway in both the cortex and hippocampus of rats. In that study, treatment with the anesthetic dexmedetomidine reduced pro-inflammatory cytokine levels and prevented inactivation of the PI3K/Akt pathway ^31^. Liu et al observed that BDNF secretion by astrocytes inhibits the pro-apoptotic anesthetic-induced depression of Akt ^26^. Mature BDNF (m-BDNF) binds at the cell surface with TrkB receptors to form a complex which activates a variety of intracellular signalling pathways, including PI3K ^32^. This is shown in Figure 3. PI3K activation phosphorylates and activates Akt in the plasma membrane. The overall effect of m-BDNF increasing activity in these pathways is inhibition of apoptosis, stimulation of new cell growth (neurones and glia) and promotion of new dendrite and axonal growth and interconnections (neuroplasticity). Liu et al suggested that anesthetic-induced neurotoxicity occurs at times of relative paucity of astrocytes and the neuroprotective factor they produce; BDNF ^26^. This paucity occurs in young and aged brains, so suggesting that adverse neurocognitive sequalae are more likely to occur when one or a combination of factors [toxins (anesthetics), inflammation (sepsis, stress, Alzheimer’s proteins, extracorporeal circulation)] come together at these vulnerable times of life when the brain natural protective mechanisms are at their weakest. The implication of this notion is clear. Using alfaxalone for the anesthetic during those times, by activating PXR will tend to bolster the natural neuroprotective mechanisms.

Two examples published in the literature support this idea. First, Yawno et al showed in fetal lambs exhibiting increased neuronal apoptosis caused by inhibition of allopregnanolone synthesis that an anesthetic dose of alfaxalone prevented the apoptosis ^33^. The second example concerns reperfusion injury in the CNS after stroke in which microglial inflammation is well established as a mechanism ^34^. Preclinical studies have shown that progesterone and allopregnanolone suppress that inflammatory response and the apoptotic enzyme caspase-3, and also decrease cerebral infarct volume after experimental brain injury^35^. Cervantes et al showed that a single anesthetic dose of alfaxalone administered immediately after restoration of normal brain perfusion prevented severe neurological damage caused by 8 minutes cardiorespiratory arrest in cats ^36^.

More recently it has been reported that alfaxalone is not neurotoxic to the developing brain ^14^. This report is reassuring but the results of the study reported herein suggest that alfaxalone anesthesia may also be accompanied by positive effects on neuronal function mediated by PXR activation, protecting neurons from the adverse effects of surgical stress and other comorbidities commonly found in patients presenting for surgery ^37–39^.

### Study Limitations

The limitation of this study lies in question of relevance of the in vitro system to the in vivo situation. The in vitro cell culture approach provides benefits of investigating h-PXR activation under controlled conditions without interference of confounding physiological or pathological factors seen in the intact animal model. However, the in vitro system is artificial and interactions between different cell types in intact animals is missing. It is therefore important to confirm in vitro findings using an intact animal model. To that end, it is important to note data already published by Yawno and colleagues that showed normal anesthetic doses of alfaxalone can replace allopregnanolone in the control of apoptosis in fetal lambs, an effect of allopregnanolone-PXR referred to above ^33^. Although the study reported herein used in vitro methods, the naturally occurring and active hormone, allopregnanolone, was used as an active control. The results show that allopregnanolone was active in this model and further that alfaxalone was more efficacious than allopregnanolone in h-PXR activation.

